# The Cl^-^-channel TMEM16A controls the generation of cochlear Ca^2+^ waves and promotes the refinement of auditory brainstem networks

**DOI:** 10.1101/2021.07.05.451099

**Authors:** Alena Maul, Saša Jovanovic, Antje K. Huebner, Nicola Strenzke, Tobias Moser, Rudolf Rübsamen, Christian A. Hübner

## Abstract

Before hearing onset (postnatal day 12 in mice), inner hair cells (IHC) spontaneously fire action potentials thereby driving pre-sensory activity in the ascending auditory pathway. The rate of IHC action potential bursts is modulated by inner supporting cells (ISC) of Kölliker’s organ through the activity of the Ca^2+^ activated Cl^-^ channel TMEM16A. Here we show that conditional deletion of *Tmem16a* in mice disrupts the generation of Ca^2+^ waves within Kölliker’s organ, reduces the burst firing activity and the frequency-selectivity of auditory brainstem neurons in the medial nucleus of the trapezoid body (MNTB), and also impairs the refinement of MNTB projections to the lateral superior olive (LSO). These results reveal the importance of the activity of Kölliker’s organ for the refinement of central auditory connectivity. In addition, our study suggests a mechanism for the generation of Ca^2+^ waves, which may also apply to other tissues expressing TMEM16A.

## Introduction

Before hearing onset around postnatal day 12 (P12) in mice (Shnerson and Pujol 1981, Sonntag, Englitz et al. 2009, Muller, Sonntag et al. 2019), the afferent auditory system exhibits cochlea-driven spontaneous activity (Lippe 1994, Jones, Leake et al. 2007). Inner hair cells (IHCs) fire bursts of action potentials (Kros, Ruppersberg et al. 1998) which drive afferent transmission to spiral ganglion neurons (SGNs) (Beutner and Moser 2001, Glowatzki and Fuchs 2002) and thus trigger bursting discharges through ascending auditory pathways (Sonntag, Englitz et al. 2009, Tritsch and Bergles 2010, Babola, Li et al. 2018). Similar to developing motor and visual systems (Katz and Shatz 1996, Hanson and Landmesser 2004), patterned activity of auditory neurons was proposed to promote activity-dependent refinement of auditory circuits before hearing onset (Clause, Kim et al. 2014, Clause, Lauer et al. 2017).

In the developing inner ear, non-sensory inner supporting cells (ISCs) form a transient epithelial structure known as Kölliker’s organ (Hinojosa 1977, Hou, Chen et al. 2019). ATP released from ISCs through connexin hemichannels (Mazzarda, D’Elia et al. 2020) activates purinergic receptors in a paracellular manner leading to cell volume decrease of ISCs and cochlear Ca^2+^ transients (Tritsch, Yi et al. 2007, Tritsch and Bergles 2010, Babola, Li et al. 2018). It was proposed that the Ca^2+^-activated Cl^-^-channel TMEM16A, which is expressed in ISCs, might be the pacemaker for spontaneous cochlear activity (Yi, Lee et al. 2013). Indeed, spontaneous osmotic cell shrinkage was shown to be mediated by TMEM16A-dependent Cl^-^ efflux, which forces K^+^ efflux from ISCs and thus the transient depolarization of IHCs (Yi, Lee et al. 2013). Thereby, bursting activity of nearby IHCs, which will later respond to similar sound frequencies, becomes synchronized (Wang, Lin et al. 2015, Eckrich, Blum et al. 2018, Harrus, Ceccato et al. 2018), establishing a possible scenario for tonotopic map refinement in central auditory structures.

Using *Tmem16a* conditional knockout mice we show that TMEM16A not only modulates ISC volume, but also drives the amplification of localized Ca^2+^ transients to propagating Ca^2+^ waves within the cochlea. Prior to hearing onset, knockout mice show reduced burst firing of neurons in the medial nucleus of the trapezoid body (MNTB), downstream of SGNs and neurons of the cochlear nucleus. Moreover, the frequency-selectivity of individual MNTB neurons is diminished shortly after hearing onset (P14) pointing towards reduced refinement of auditory connections. Indeed, neurons from the lateral superior olive (LSO) received twice as many functional MNTB afferents in knockout mice compared to wildtype littermates. Taken together these results suggest that the Ca^2+^-activated Cl^-^-channel TMEM16A plays a significant role in Ca^2+^ wave generation and contributes to the refinement of auditory brainstem circuitries prior to hearing onset.

## Results

### TMEM16A is required for the generation of cochlear Ca^2+^ waves

The Ca^2+^ activated Cl^-^ channel TMEM16A is expressed in ISCs of Kölliker’s organ (for a schematic representation of the organ of Corti see Figure 1A) (Yi, Lee et al. 2013, Wang, Lin et al. 2015), and is activated by ATP-induced increase in Ca^2+^ concentration (Yi, Lee et al. 2013, Wang, Lin et al. 2015). The opening of TMEM16A triggers Cl^-^ efflux, followed by K^+^ efflux and cell shrinkage. The ensuing rise of extracellular K^+^ drives electrical activity of immature IHCs (Wang, Lin et al. 2015). To assess the role of TMEM16A in the developing cochlea and the impact of TMEM16A-dependent cochlear signaling on the development of auditory brainstem nuclei we disrupted *Tmem16a* in the inner ear. This was achieved by mating our floxed *Tmem16a* line (*Tmem16a^fl/fl^*) (Heinze, Seniuk et al. 2013) with a line expressing Cre-recombinase under the control of the *Pax2* promoter (Ohyama and Groves 2004), which is active in the otic placode (Lawoko-Kerali, Rivolta et al. 2002). This line is subsequently referred to as cKO mice. *Tmem16a* deletion was confirmed by immunohistochemistry (Figure S1A) and Western blot analysis (Figure S1B). Importantly, organs of Corti of cKO mice showed no obvious morphological defects. The development of the tectorial membrane and the morphology of the hair cells appeared normal before hearing onset (P6), at hearing onset (P12) or in the weeks thereafter (3 weeks and 6 weeks after birth) (Figure S1C and D) indicating that TMEM16A and TMEM16A-dependent activity of Kölliker’s organ is not essential for the morphologic development of the organ of Corti.

**Figure 1.**
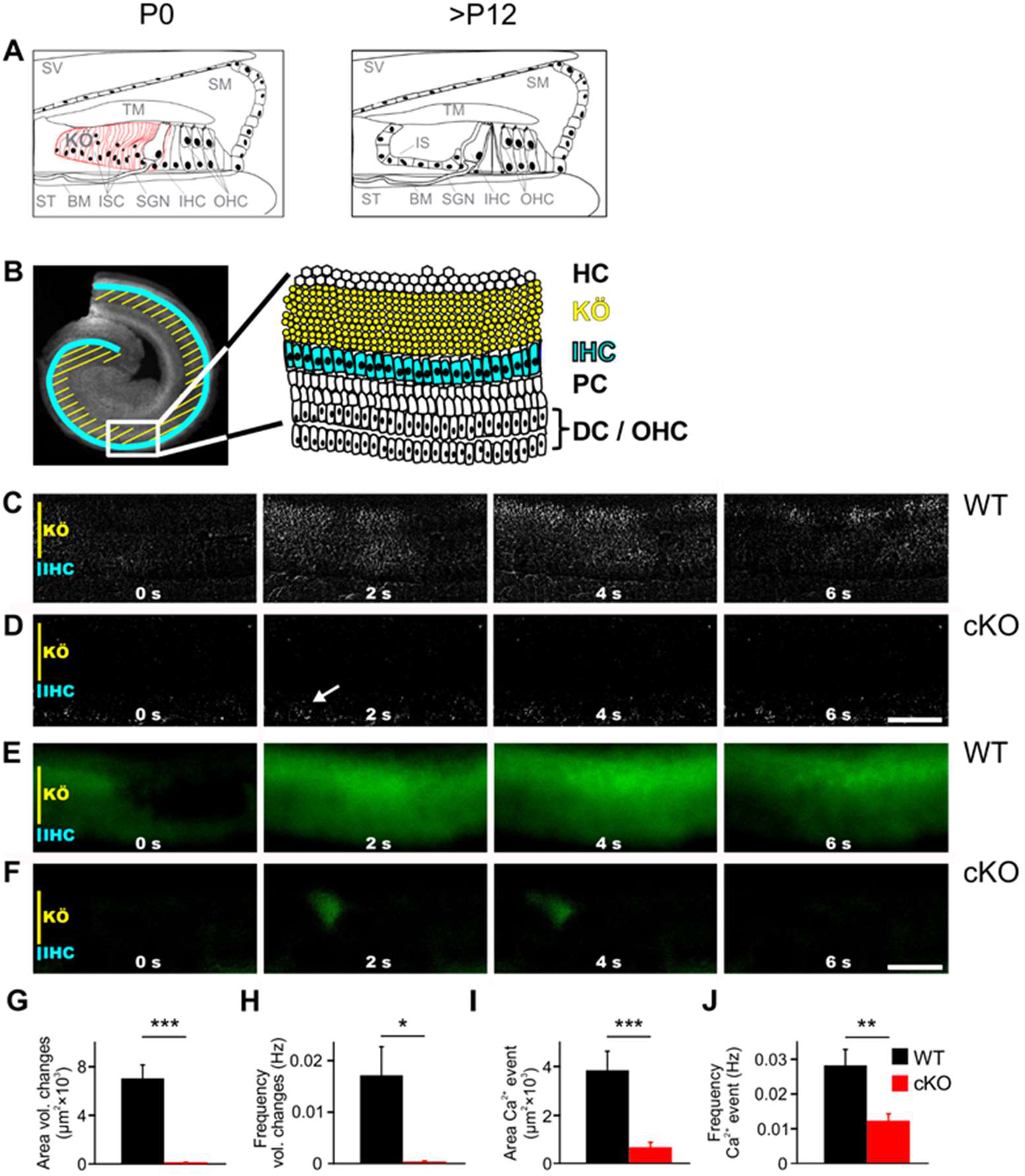
TMEM16A is required for the generation of spontaneous volume changes and Ca^2+^ waves in Kölliker’s organ. (**A**) Schematic representation of the organ of Corti at birth (left) and after hearing onset (right). BM, basilar membrane; IHC, inner hair cells; IS, inner sulcus; ISC, inner supporting cells; KÖ, Kölliker’s organ; OHC, outer hair cells; SGN, spiral ganglion neurons; SM, scala media; ST, scala tympani; SV, scala vestibuli; TM, tectorial membrane. (**B**) Top view showing an isolated apical cochlea turn and the location of inner hair cells (IHCs) (*blue*) and Kölliker’s organ (KÖ) (*yellow*). The zoom-in shows a schematic drawing of the area imaged in C-F. DC, Deiter’s cells; HC, Hensen’s cells; OHC, outer hair cells; PC, pillar cells. (**C, D**) DIC timelapse imaging at P7 reveals spontaneous volume changes of inner supporting cells (ISCs) in a wildtype mouse indicated by changes in light intensity, which are almost absent in the cKO littermate. The arrow in D indicates erythrocytes moving in a blood vessel. Scale bar 50 μm. (**E, F**) Ca^2+^ imaging at P6 reveals spontaneous Ca^2+^ waves traveling across ISCs of Kölliker’s organ in a wildtype mouse (E) that are reduced to small local Ca^2+^ transients in the cKO littermate (F). Scale bar 50 μm. (**G, H**) Quantification of areas and frequencies of spontaneous ISC volume changes. Values represent mean ± SEM (P5-7; n=7 WT, n=9 cKO; two-tailed unpaired Student’s t-test: area: p<0.001, frequency: p<0.05). (**I, J**) Quantification of area and frequency of spontaneous Ca^2+^ events. Values represent mean ± SEM (P5-7; n=14 WT, n=16 cKO; two-tailed unpaired Student’s t-test: area: p<0.001; frequency: p<0.01).

To investigate the volume changes and Ca^2+^ waves that spontaneously appear in the ISCs of Kölliker’s organ (Tritsch, Yi et al. 2007, Anselmi, Hernandez et al. 2008, Tritsch and Bergles 2010), acutely isolated cochleae from P5-7 wildtype and cKO littermates were used (Figure 1B). While wildtype cochleae showed notable volume changes of groups of ISCs (Figure 1C, G, H; n=7; mean event area ± SEM= 7020 ± 1130 μm^2^; mean event frequency ± SEM= 0.0171 ± 0.0056 Hz), volume changes were almost absent in cochleae acutely isolated from 5-7-day-old cKO mice (Figure 1D, G, H; n=9; mean event area ± SEM= 40 ± 30 μm^2^, p<0.001; mean event frequency ± SEM= 0.0003 ± 0.0002 Hz, p<0.05), in agreement with previously published results (Wang, Lin et al. 2015). In wildtype cochleae, these events propagated in waves along the tonotopic axis of the cochlea also affecting phalangeal cells that surround IHCs (Video S1). To visualize changes in intracellular Ca^2+^ concentrations, cochlear explants were loaded with the Ca^2+^ indicator dye Fura-2 AM. Wildtype mice showed Ca^2+^ transients which travelled in waves along the tonotopic axis of Kölliker’s organ (Figure 1E, I, J; n=14; mean event frequency ± SEM= 0.0282 ± 0.0046 Hz; mean event area ± SEM= 3840 ± 790 μm^2^). In cochleae from cKO littermates, Ca^2+^ transients were rare and restricted to small areas (Figure 1F, I, J, Video S2; n=16; mean event frequency ± SEM= 0.0123 ± 0.0020 Hz; p<0.01; mean event area ± SEM= 660 ± 210 μm^2^; p<0.001).

Spontaneous Ca^2+^ waves are elicited by ATP-induced ATP release from ISCs (Mazzarda, D’Elia et al. 2020), a mechanism possibly involving P2Y1, P2Y2 and P2Y4 receptors (Piazza, Ciubotaru et al. 2007, Babola, Li et al. 2018), Cx26 and Cx30 heteromeric hemichannels (Anselmi, Hernandez et al. 2008, Schutz, Scimemi et al. 2010) and TMEM16A (Wang, Lin et al. 2015). To assess what role TMEM16A might play in ATP-mediated Ca^2+^ signals we applied the non-selective P2 receptor antagonist suramin (150 μM) (von Kugelgen and Wetter 2000, Burnstock 2014). In wildtype mice, suramin application reduced Ca^2+^ waves to uncoordinated, locally restricted Ca^2+^ transients (n=6; mean event area before suramin application ± SEM= 4476 ± 1007 μm^2^ and after suramin application ± SEM= 1172 ± 389 μm^2^; p<0.05 paired two-tailed Student’s t-test). Suramin application had no effect on Ca^2+^ transients in cKO mice (n=5; mean event area before suramin application ± SEM= 466 ± 187.8 μm^2^ and after suramin application ± SEM= 378 ± 112 μm^2^; p=0.24 paired two-tailed Student’s t-test) (Figure S2). This supports the notion that TMEM16A drives the ATP release from ISCs and is important for the spreading of spontaneous activity between ISCs of Kölliker’s organ via P2 receptors.

### TMEM16A-dependent cochlear activity modulates the burst firing pattern of MNTB neurons

By increasing the K^+^ concentration, TMEM16A-dependent activity of Kölliker’s organ leads to the generation of Ca^2+^ action potentials in IHCs. This is followed by Ca^2+^ dependent exocytosis of glutamate at the IHC synapse, which drives burst firing of action potentials in SGNs (Wang, Lin et al. 2015). The bursting activity is then relayed to central auditory neurons (Tritsch and Bergles 2010, Babola, Li et al. 2018) (Figure 2A shows a schematic representation of the auditory brainstem), and is believed to be important for the proper development of synaptic contacts and tonotopic maps (Clause, Kim et al. 2014, Clause, Lauer et al. 2017).

**Figure 2.**
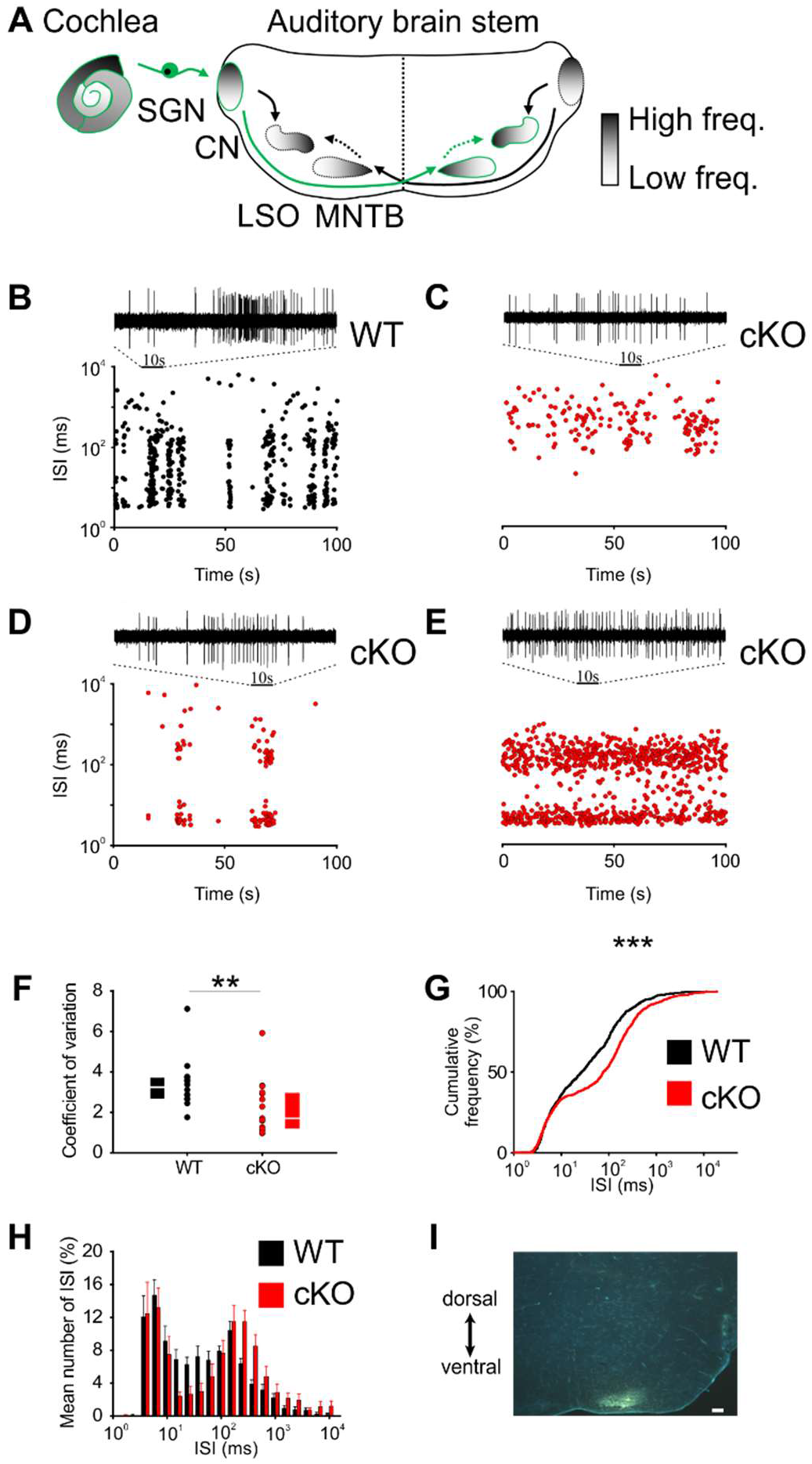
Disruption of Kölliker’s organ activity changes prehearing burst firing of MNTB neurons *in vivo*. (**A**) Simplified model of auditory connections in the brainstem. The pathways relevant for this experiment are marked in green. Inhibitory pathways are indicated by dotted arrows. CN, cochlear nucleus; MNTB, medial nucleus of the trapezoid body; LSO, lateral superior olive; SGN, spiral ganglion neurons. (**B - E**) Patterns of spontaneous discharge activity from individual MNTB neurons recorded from mice before hearing onset (P8) in wildtype and cKO littermates. Dot-plot graphs show respective interspike interval (ISI) distributions for 100 s of spontaneous discharge activity. On top of each dot-plot raster is a 10 s period of original spike trains. Note that the wildtype MNTB neuron shows prominent burst firing, which is either strongly reduced (D) or absent in cKO mice (C and E). (**F**) Quantification of spike bursting patterns by calculating the coefficient of variation of ISIs yields significant differences between wildtype (n=14) and cKO units (n=15) (Mann-Whitney rank sum test: p<0.01); also shown are box-plots indicating medians and 25 % and 75 % quartiles. (**G**) The mean cumulative distribution of ISIs reveals the significant shift towards larger values in cKO mice (wildtype n=14 and cKO n=15; Kolmogorov-Smirnov test: p<0.001, D=0.19), with the median ISI increasing from 26.9 ms in wildtype to 76.3 ms in cKO mice. (**H**) The overlaid log-binned histogram compares the distribution of ISIs between wildtype and cKO mice. Values represent mean ± SEM (n=14 wildtype, n=15 cKO (P8)). For statistical analysis, the Chi Square test was used (see Table S1 for p-values). (**I**) Iontophoretic injection with Fluorogold verifies recording site from *in vivo* juxtacellular voltage recordings from MNTB neurons in a prehearing wildtype mouse (P8). Scale bar 200 μm.

To investigate effects of *Tmem16a* knock-out on the burst firing patterns in auditory brainstem neurons, juxtacellular single unit recordings from MNTB neurons were obtained from *in vivo* prehearing mice (P8). Similar to SGNs (Tritsch, Rodriguez-Contreras et al. 2010, Wong, Jing et al. 2013), MNTB neurons exhibit spontaneous bursts of spikes that could last several seconds (Figure S3A) (Sonntag, Englitz et al. 2009). These long bursts are made up of a series of short “mini-bursts”, consisting of several spikes, and occurring at intervals of approx. 100 ms (Fig. S3B, C).

Bursting activity was quantified by the coefficient of variation of interspike intervals (CV), whereby values below 1 correspond to random firing and higher values indicate more patterned activity (Jones, Leake et al. 2007). In wildtype mice, MNTB neurons showed the typical bursting activity (Figure 2B,F; n=14, median CV [25%, 75% quartiles]= 3.2 [2.65, 3.56]) (Sonntag, Englitz et al. 2009), while in cKO littermates MNTB firing patterns had significantly smaller CVs (Figure 2C-F; n=15, median CV [25%, 75% quartiles]= 1.7 [1.2, 2.9]; p<0.01, Mann-Whitney test). Intermediate interspike intervals (ISI), representing the interval between mini-bursts, were severely diminished in all MNTB neurons recorded from cKO mice. A presentation of the firing pattern as mean cumulative distribution of ISIs (Figure 2G; Kolmogorov-Smirnov test: p<0.001, D=0.19) and in an overlaid log-binned histogram (Figure 2H; Chi square test, p-values are shown in Table S1) illustrates a shift towards longer ISIs in cKO mice. The additional detailed analysis of burst firing patterns in MNTB neurons revealed further significant differences between wildtype and cKO littermates for (i) the number of bursts per 100 s (median [25%, 75% quartiles]: WT= 3.9 [3.3, 5.8]; cKO= 0.6, [0, 3.8]; p<0.01, Mann-Whitney test), (ii) the duration of bursts (median [25%, 75% quartiles]: WT= 1.6 [0.9, 3.3]; cKO= 0.7 [0.3, 1.8]; p<0.001, Mann-Whitney test), and (iii) the number of spikes per burst (median [25%, 75% quartiles]: WT= 45 [21, 89]; cKO= 19 [14, 31]; p<0.001, Mann-Whitney test) (Figure 3A-C). MNTB neurons recorded from cKO mice showed a large variation of action potential firing rates, but still the mean firing rate did not differ from wildtype cells (Figure 3D; mean firing rate ± SEM: WT= 6.2 ± 0.8 Hz, n=14; cKO= 5.8 ± 1.1 Hz, n=15; p=0.69, Student’s t-test).

**Figure 3.**
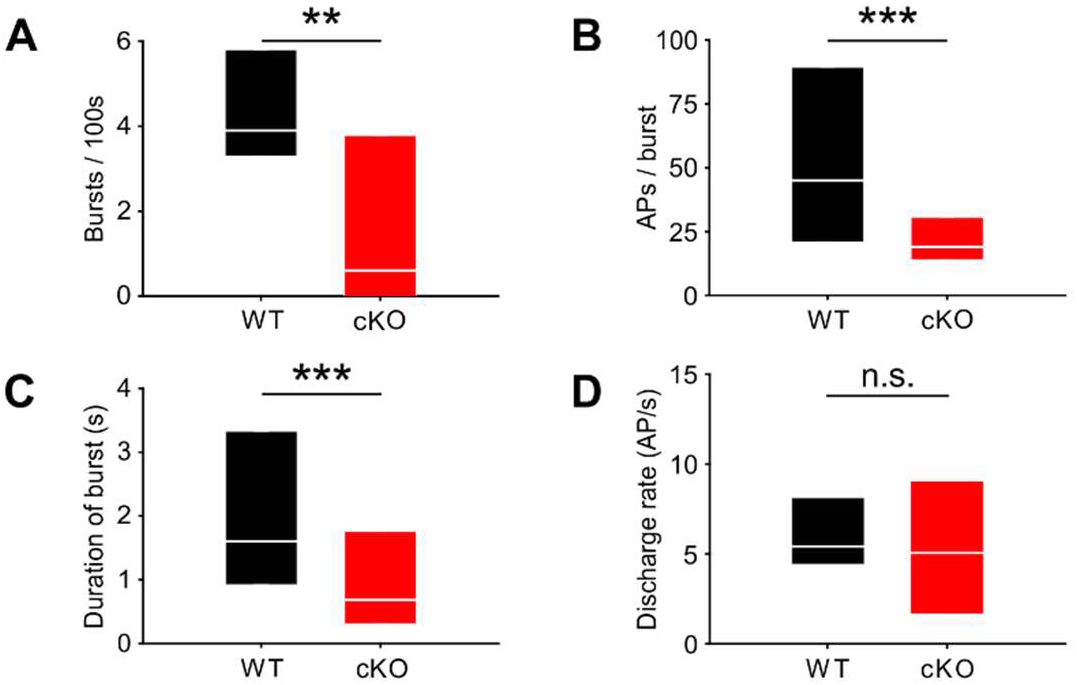
Lack of TMEM16A in cochlear ISCs changes bursts but not the overall firing rate in MNTB auditory brainstem neurons *in vivo*. (**A - C**) Spontaneous discharge patterns of MNTB neurons recorded from cKO mice show a reduced number of bursts per 100 s (A), a reduced number of spikes per burst (B) and a reduced duration of bursts (C) compared to wildtype. Values represent median with 25 %, 75 % quartiles (n=14 WT; n=15 cKO (P8); Mann-Whitney rank sum test: Number of bursts per 100 s: p=0.01; Number of spikes per burst: p<0.001; duration of burst: p<0.001). (**D**) The overall discharge rates did not differ between wildtype (n=14) and cKO (n=15) (two-tailed unpaired Student’s t-test: p=0.69).

Since TMEM16A is neither expressed in SGNs nor in CN, MNTB, and LSO neurons in wildtype mice (Figure S4) and expression in the brainstem was limited to vascular smooth muscle cells, we primarily attribute the severely altered burst firing activity to impaired Kölliker’s organ activity in cKO mice.

### Frequency-selectivity of MNTB neurons is reduced in cKO mice

To test whether the changes in burst firing patterns of cKO MNTB neurons have consequences on neuronal function after hearing onset, auditory brainstem responses (ABRs) were measured at P13-14. cKO mice had similar ABR thresholds in response to stimulation with clicks or tone-bursts at 6 kHz, 12 kHz, and 24 kHz as wildtypes (Figure 4A,B, Table S2; n=6 WT; n=7 cKO). In both genotypes, ABRs to click stimuli of various intensities (40–100 dB) mainly consisted of three waves (labelled I-III) which were comparable in latency and amplitude (Figure S5, Table S3, S4). These data indicate that cKO mice have a normal sensitivity to sound stimulation and normal temporal precision of the spiking response to sound onset in the lower auditory pathway.

**Figure 4.**
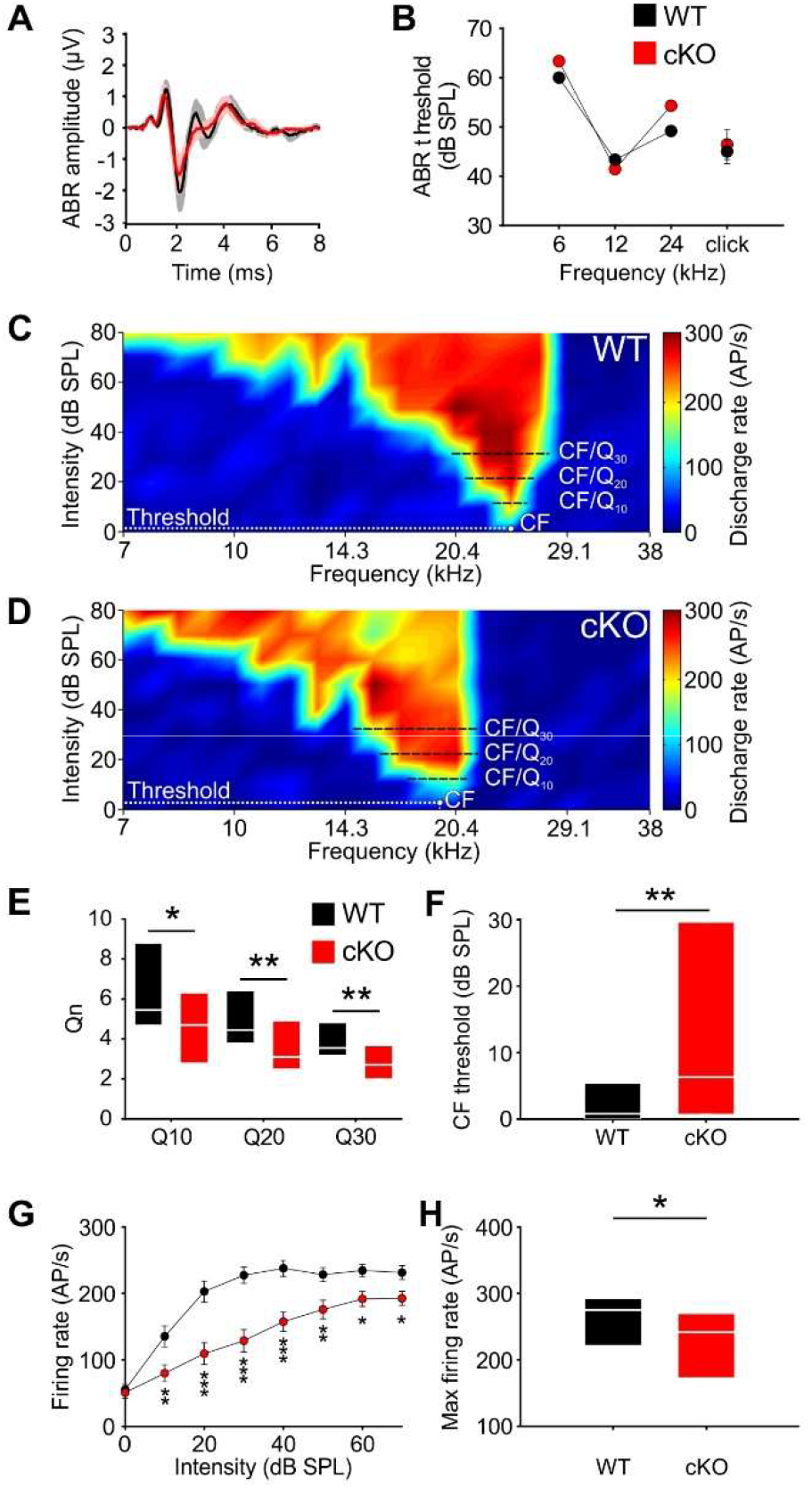
Similar ABR thresholds, but differences in frequency-selectivity, response threshold and maximal firing rate of MNTB neurons in cKO mice. (**A**) Grand averages of ABR waveforms to 80 dB click stimulation recorded from cKO (red) and wildtype (black) at P13-14. (**B**) ABR thresholds in response to 6, 12 or 24 kHz tone bursts and click stimuli did not differ between wildtype (n=7) and cKO (n=6) at P13-14. For values, see Table S2. (**C, D**) Representative frequency response areas of MNTB neurons (P14) recorded juxtacellularly in a wildtype mouse (C) (CF: 24 kHz, threshold 0.1 dB SPL, Q10/20/30 = 6.4/4.3/4.2), and in a cKO littermate (D) (CF: 18.4 kHz, threshold= 3 dB SPL, Q10/20/30= 3.7/3.2/2.8). Note that the response area is broader and that the CF thresholds are increased in cKO. (**E**) Frequency-selectivity of MNTB neurons was reduced in cKO mice as indicated by lower Q-factors (shown are medians and 25 %, 75 % quartiles (n=20 WT; n=25 cKO (P14); Mann-Whitney rank sum test: Q10: p<0.05, Q20: p<0.01, Q30: p<0.01). (**F**) Sound thresholds of individual MNTB neurons are elevated in cKO mice compared to wildtype (p<0.01). (**G**) Average rate level functions at characteristic frequency show decreased action potential firing in cKO at SPLs above 10 dB (two-way ANOVA: effect of strain p<0.001, effect of intensity p<0.001). (**H**) Maximum firing rates during acoustic stimulation are significantly decreased in cKO mice compared to wildtype littermates (two-tailed unpaired Student’s t-test: p<0.05).

Next, we assessed whether the disruption of TMEM16A-dependent cochlear activity affects the frequency tuning properties of MNTB neurons. Therefore, the frequency response areas (FRAs) of single MNTB neurons were acquired in cKO and wildtype littermates using *in vivo* electrophysiology and tone burst stimulation. Juxtacellular recordings were performed at P14, i.e. shortly after the onset of hearing to avoid possible compensatory effects of acoustically driven activity on neuron responsiveness (Werthat, Alexandrova et al. 2008, Bogart, Levy et al. 2011). The characteristic frequencies (CF) of the recorded MNTB neurons, i.e., the frequency value at which the neuron is excited with the minimal intensity possible, ranged between 5.3 and 30.5 kHz (mean CF ±SEM: WT= 15.4 ±1.1 kHz (n=32); cKO= 16.2 ±1.3 kHz (n=25)). MNTB neurons recorded from wildtype mice showed the typical V-shaped FRAs (Sonntag, Englitz et al. 2009) with acoustically driven excitation sharply narrowing towards lower intensities (Figure 4C). The filter characteristics of the FRAs were quantified by the Q_n_-value, a measure of the unit’s sharpness of tuning, which is calculated as the ratio of CF to bandwidth at 10, 20 and 30 dB above threshold (e.g. Q_10_=CF/BW10 with BW_10_=bandwidth at 10 dB above threshold). For wildtype mice the median [25%, 75% quartiles] was Q_10_= 5.5 [4.7, 8.8], Q_20_= 4.5 [3.8, 6.4], and Q_30_= 3.6 [3.2, 4.8]) (Figure 4E). Neurons recorded from the cKO littermates (n=25) had significantly broader excitatory response areas, i.e. lower frequency selectivity as indicated by significantly lower Q-factors at all three above-threshold levels (Figure 4D,E; median [25%, 75% quartiles]: Q_10_= 4.7 [2.8, 6.3], p<0.05; Q_20_= 3.1 [2.5, 4.9], p<0.01; Q_30_= 2.7 [2.0, 3.7], p<0.01, Mann-Whitney test). Additionally, MNTB neurons in cKO mice had elevated thresholds in comparison to the wildtype littermates (median threshold [25%, 75% quartiles]: 0.8 [0.0, 5.4] dB SPL) (Figure 4F; median [25%, 75% quartiles]: 6.3 [0.8, 29.6] dB SPL; p<0.01, Mann-Whitney test). Overall CF threshold levels tended to show a larger variability in knockout compared to wildtype mice (cKO: −7.3 dB SPL to 48.3 dB SPL; WT: −8.4 dB SPL to 18.7 dB SPL). Furthermore, the sound-evoked firing properties of the MNTB neurons in cKO mice were also affected. The rate-level functions at CF showed significantly lower firing rates for sound intensities at and above 10 dB SPL of toneburst stimulation in comparison to wildtype littermates (effect of strain p<0.001, effect of intensity p<0.001, interaction strain × intensity p=0.006; two-way ANOVA) (Figure 4G). Maximal firing rate of individual neurons in response to any CF/intensity combination was markedly diminished in cKO (mean FR ± SEM: WT= 263.6 ± 12.4 action potentials/s, n=25; cKO= 223.9 ± 11.1 action potentials/s, n=32; p<0.05, t-test) (Figure 4H). Apparently, MNTB neurons in cKO mice cannot respond to high intensity stimulation with sufficient firing rates that are typically observed in wildtype littermates. Taken together, these data demonstrate that frequency selectivity and sensitivity to acoustic stimulation in single MNTB neurons are impaired upon disruption of *Tmem16a* in the cochlea.

### The developmental refinement of functional connections of the MNTB-LSO pathway is impaired in cKO mice

Despite the above-described differences in the pattern of spontaneous and sound-evoked activity in MNTB neurons between wildtype and cKO mice, the gross morphology of the MNTB and the LSO appeared normal (mean MNTB area ± SEM: WT= 63×10^3^ ± 4.6 μm^2^ (n=10); KO= 55×10^3^ ± 3.4 μm^2^ (n=10); p=0.08; mean LSO area ± SEM: WT= 117×10^3^ ± 6.8 μm^2^ (n=10); KO= 110×10^3^ ± 3.9 μm^2^ (n=10); p=0.43).

Since spontaneous burst activity of auditory neurons might promote the targeting and refinement of their projections as suggested for the developing visual system (Torborg, Hansen et al. 2005), we assessed the synaptic and topographic refinement of MNTB and LSO neurons. MNTB neurons were activated via photolysis of caged glutamate and the number of functional inputs on individual LSO neurons was assessed by whole-cell current-clamp recordings. In a pilot experiment the distance was measured at which glutamate uncaging produces action potentials in MNTB neurons. The mean distance (± SEM) was 19.2 ± 0.8 μm mediolaterally and 20.0 ± 1.3 μm dorsoventrally indicating that glutamate uncaging was locally restricted to a surface area of 20×20 μm^2^ and that light scattering that might influence the amount of uncaged glutamate in the tissue was negligible (Figure S6A-C).

Notably, the size of MNTB input areas in cKO mice was doubled compared to wildtype (Figure 5A-C; mean cross-sectional input area ± SEM: WT= 10 ± 1 % (n=10) and cKO: 20 ± 2 % (n=10) of the respective MNTB cross-sectional areas; p<0.01, Student’s t-test). Moreover, the input width, defined as the maximal distance of responsive uncaging sites along the mediolateral axis, was twice as large (Figure 5A, B, D; mean input width ± SEM: WT= 18 ± 3 % of the MNTB’s mediolateral length; cKO= 36 ± 5 % of the MNTB’s mediolateral length; p<0.01, Student’s t-test). The large increase in MNTB input width in cKO mice reveals that LSO neurons located in the medial high frequency region of the nucleus not only received input from neurons of the high frequency (medial) region of the MNTB, but also from neurons in the mid-nuclear region tuned to lower frequencies. For additional examples comparing the size and width of MNTB input areas between wildtype and cKO mice see Figure S5D,E. These data strongly point towards an impairment of the tonotopic refinement of MNTB-to-LSO projections in cKO mice.

**Figure 5.**
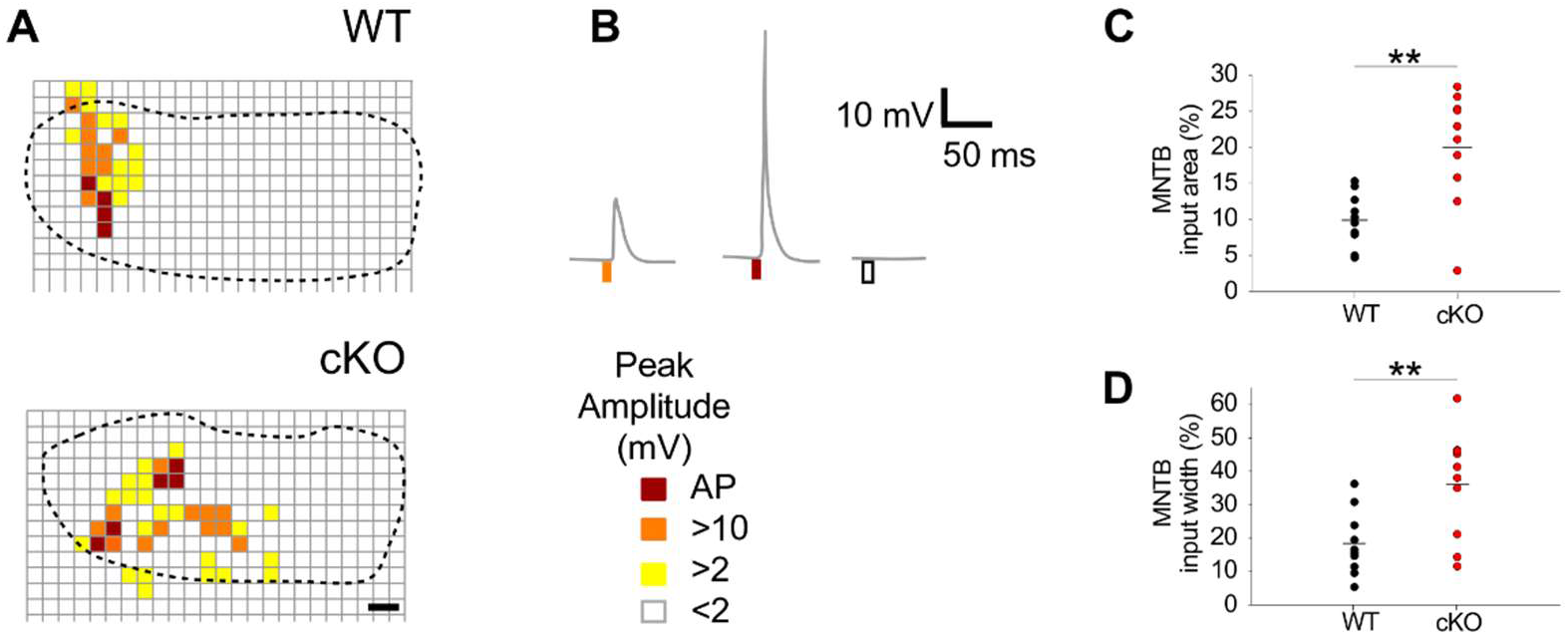
MTNB-LSO input maps are enlarged upon disruption of TMEM16A. (**A**) Exemplary MNTB input maps from wildtype (P9) and cKO mice (P10) as revealed by whole-cell current clamp recordings (*dotted line* outlines the MNTB area, *grid points* indicate glutamate uncaging sites). The location of responsive (*colored squares*) and unresponsive (*open squares*) uncaging sites are indicated. Scale bar 40 μm. (**B**) Uncaging of glutamate close to presynaptic MNTB neurons gives rise to synaptic responses of various peak amplitudes (*left*), elicits action potentials (*middle*), or fails to evoke significant voltage signals (*right*) in the recorded LSO neurons (see color code below). (**C** and **D**) Quantification of MNTB input areas (summed area of all responsive uncaging sites) on a recorded LSO neuron and of the MNTB input widths (maximal distance of stimulations sites that evoked depolarization greater than 10 mV along the mediolateral (tonotopic) axis of the MNTB). MNTB input areas were normalized to the corresponding MNTB cross-sectional area and MNTB input widths to the length of the mediolateral axis of the MNTB. Values represent mean ± SEM (n=10 WT; n=10 cKO (P9-11); two-way unpaired Student’s t-test: MNTB input area: p<0.01, MTNB input width: p<0.01).

## Discussion

Kölliker’s organ was identified as the origin of pre-sensory cochlear activity, which was suggested to serve the refinement of auditory circuits (Tritsch, Yi et al. 2007). The Ca^2+^-activated Cl^-^-channel TMEM16A, which is expressed in Kölliker’s organ shortly before hearing onset, mediates spontaneous osmotic cell shrinkage of ISCs, which forces K^+^ efflux and thus the transient depolarization of IHCs (Yi, Lee et al. 2013, Wang, Lin et al. 2015). Thereby, bursting activity of nearby IHCs, which will later respond to similar sound frequencies, becomes synchronized (Wang, Lin et al. 2015, Eckrich, Blum et al. 2018, Harrus, Ceccato et al. 2018), establishing a possible scenario for tonotopic map refinement (Tritsch, Yi et al. 2007, Tritsch and Bergles 2010, Johnson, Eckrich et al. 2011). Direct proof for a correlation between TMEM16A-mediated ATP release and tonotopic map refinement remained elusive. Here, we demonstrate that disruption of TMEM16A in the inner ear impairs prehearing cochlear activity and impairs spontaneous burst firing as well as sensitivity and frequency selectivity of sound-evoked firing of MNTB neurons. Finally, we show that these alterations in prehearing auditory activity severely impair the refinement of the MNTB-LSO pathway.

Spontaneous activity of Kölliker’s organ is characterized by recurrent ATP-dependent changes in cell volume and Ca^2+^ transients of ISCs (Tritsch, Yi et al. 2007). The decreases in cell volume reflect the passive movement of water associated with Cl^-^ efflux upon activation of the Ca^2+^-activated Cl^-^-channel TMEM16A (Yi, Lee et al. 2013, Wang, Lin et al. 2015). Since the volume changes of ISCs coincided with the generation of cochlear Ca^2+^ waves, we hypothesized that TMEM16A may play a role for the propagation of Ca^2+^ events. Indeed, Ca^2+^ imaging of acutely isolated cochleae revealed that Ca^2+^ waves were reduced to isolated local Ca^2+^ events in cKO mice. Since suramin blocked Ca^2+^ waves in wildtype cochleae but had no effect on local Ca^2+^ transients in cKO mice, the effect of TMEM16A on the propagation of Ca^2+^ transients along Kölliker’s organ clearly involves the activation of purinergic receptors. ATP release by ISCs seems to occur through the opening of connexin hemichannels triggered by Cl^-^ efflux-mediated osmotic cell shrinkage (Mazzarda, D’Elia et al. 2020). Given the presence of mechanically induced ATP release and purinergic receptor activation in the generation of Ca^2+^ waves in other tissues expressing TMEM16A such as the airway and submandibular gland epithelium (Sanderson, Charles et al. 1990, Felix, Woodruff et al. 1996, Grygorczyk and Hanrahan 1997, Watt, Lazarowski et al. 1998, Homolya, Watt et al. 1999, Homolya, Steinberg et al. 2000, Romanenko, Catalan et al. 2010, Ryu, Peixoto et al. 2010), the olfactory epithelium (Hegg, Irwin et al. 2009, Henriques, Agostinelli et al. 2019), and the biliary epithelium (Dutta, Woo et al. 2013), it will be interesting to investigate whether ATP/TMEM16A-mediated propagation of Ca^2+^ waves is a wide-spread feature in many tissues.

To analyze the impact of the disruption of TMEM16A on the auditory pathway we performed *in vivo* juxtacellular recordings in the MNTB of cKO mice before and shortly after hearing onset. Burst firing pattern of MNTB neurons was often completely absent or severely diminished in P9 cKO mice. MNTB neurons from P14 cKO mice also showed markedly diminished maximal firing rates indicating that they cannot respond to high intensity stimulation with sufficient firing rates that are normally observed in wildtype littermates. As high sound intensities are coded by high firing rates, interaural level differences might be affected and hence also sound source localization in cKO mice. Furthermore, individual MNTB neurons recorded from P14 cKO mice responded to sound stimuli from a much larger frequency range compared to wildtype mice. The reduced frequency-selectivity could result from superfluous functional projections from cochlear onto MNTB neurons, which is likely a consequence of reduced bursting activity earlier in development. To test whether projection patterns between auditory nuclei in cKO mice are indeed altered, we examined MNTB projections to the LSO, a part of the sound localization pathway in which refinement has been well studied (Kim and Kandler 2003, Clause, Kim et al. 2014, Muller, Sonntag et al. 2019). The necessary spatial resolution was achieved with a fast galvanometric photolysis system which allowed a 5-fold improvement in area resolution compared to fiber optic-based uncaging systems used in previous studies (Kim and Kandler 2003, Kim and Kandler 2010, Clause, Kim et al. 2014). In cKO mice the number of functional connections of single LSO afferents (MNTB input area) and the respective mediolaterally covered area, i.e., the area along the tonotopic axis (MNTB input width), was twice as large as in wildtype. Since it has been reported that LSO neurons normally lose about 50 % of their afferents during the first two weeks of postnatal development of the mouse (Noh, Seal et al. 2010), the present results imply that silencing of synaptic connections was strongly diminished in cKO mice.

Lack of expression of TMEM16A in SGNs and auditory brainstem nuclei suggests that curtailed bursting activity and the paucity of tonotopic refinement of auditory projections within the auditory brainstem are caused by diminished prehearing cochlear activity that likely stems from the failed synchronization of IHC acitivity in cKO mice. Namely, if Cl^-^ efflux through TMEM16A is absent in cKO mice, simultaneous K^+^ release would not be triggered, and hence, the firing patterns of neighboring IHCs would not be synchronized (Wang, Lin et al. 2015).

This assumption is in agreement with the recent discovery that synchronization of Ca^2+^ signals of neighbouring OHCs is required for proper refinement of afferent projections onto OHCs as well as the maturation of OHCs (Ceriani, Hendry et al. 2019). The synchronization of OHC activity is also mediated by ATP release from supporting cells, namely Deiter’s cells, during the first two postnatal weeks in the mouse (Jeng, Ceriani et al. 2020).

Besides spontaneous activity of ISCs, IHC firing patterns are also modulated by cholinergic medial olivocochlear efferents (Glowatzki and Fuchs 2000, Johnson, Eckrich et al. 2011, Sendin, Bourien et al. 2013), which transiently innervate immature IHCs (Warr and Guinan 1979, Simmons, Mansdorf et al. 1996a, Simmons, Moulding et al. 1996b). In α9 knockout mice this cholinergic input is disrupted (Clause, Kim et al. 2014). The fact that α9 knockout mice show more subtle changes in the bursting behavior of MNTB neurons *in vivo* (Clause, Kim et al. 2014) than our cKO mice suggests a prevalent role of the activity of ISCs in shaping firing patterns of the early auditory pathway prior to onset of hearing. Still, our results showed reduced synaptic refinement before hearing onset similar to those of α9 knockout mice (Clause, Kim et al. 2014), and to otoferlin knockout mice, which exhibit drastically diminished spontaneous activity (Muller, Sonntag et al. 2019) due to disrupted glutamate release from IHCs (Roux, Safieddine et al. 2006). Thus, it seems that both subtle and drastic changes in the temporal pattern and/or the number of spikes lead to a severe disruption of tonotopic maps. These changes have permanent consequences, since tonotopic refinement of MNTB-LSO projections does not take place after hearing onset (Walcher, Hassfurth et al. 2011, Muller, Sonntag et al. 2019). For example, the above-mentioned circuit anomalies observed in α9 knockout mice led to diminished sound localization and sound frequency processing after hearing onset as revealed by behavioral studies (Clause, Lauer et al. 2017).

Whether TMEM16A-mediated prehearing activity has a similar impact on hearing after hearing onset remains to be determined. While ABR measurements in our study did not reveal gross alterations in signaling along the lower auditory pathway, our recordings from individual auditory brainstem neurons showed significantly elevated sound thresholds and diminished frequency selectivity in cKO mice after hearing onset (P14), indicating impaired auditory processing. This is supported by findings showing that mutations in ion channels (hemichannel, gap junctions, and Ca^2+^ channels) involved in prehearing activity of Kölliker’s organ are linked to congenital deafness (Majumder, Crispino et al. 2010).

## Experimental Procedures

### Mouse strains

Mouse care and usage were in accordance with the German animal protection laws and were approved by the local authorities. *Tmem16a^fl/fl^* mice were crossed with a *Pax2^Cre^* mouse line (a gift from A. Groves, House Ear Institute, Los Angeles) to obtain *Tmem16a* ear conditional knockout mice. Throughout the paper, these mice are referred to as cKO mice. Information about the mouse strains including genotyping primers can be found in Heinze *et al*. 2013 (Heinze, Seniuk et al. 2013) and Ohyama & Groves 2004 (Ohyama and Groves 2004). TMEM16B knockout mice were a gift from T. Jentsch, Leibniz Institute for Molecular Pharmacology (Germany) and were described in Billig et al. 2011 (Billig, Pal et al. 2011).

### Immunohistochemistry

Cochlear cryosections were prepared from mice at postnatal ages P0, P3, P9, P13 and P15. Mice were decapitated and inner ears (cochlea capsule attached to the vestibular apparatus) were isolated and processed according to a protocol from Whitlon et al. 2001 (Whitlon, Szakaly et al. 2001). In brief, inner ears were fixed at room temperature (RT) in 4 % paraformaldehyde (PFA) in phosphate-buffered saline (PBS) for 1-1.5 h. Inner ears from mice older than P5 were decalcified in 10 % EDTA (PBS) at 4°C for 12 h to up to 48 h depending on the age of the mouse. Inner ears were cryoprotected with solutions with increasing sucrose concentrations (10-30 % sucrose (PBS)) and kept in tissue Tek (Sakura) at 4 °C. Embedded inner ears were shock frozen using a dry ice / ethanol bath. Cryosections (10 μm) of inner ears were cut in the coronal plane using a cryostat (Leica CM-3050S cryostat). Whole mount brain slices were prepared from mice at P8. Coronal slices (150 μm) were cut on a vibrating microtome (Leica VT1200S) in dissection solution containing (in mM): 50 NaCl, 2.5 KCl, 1.25 NaH_2_PO_4_, 3 MgSO_4_, 0.1 CaCl_2_, 75 sucrose, 25 D-glucose, 25 NaHCO_3_ and 2 Na-pyruvate (280 mosmol, pH 7.2).

Cochlea cryosections and brain slices were fixed in 4 % PFA in PBS at RT for 20 min and incubated in blocking solution containing 1 % fetal bovine serum (BSA), 3 % goat serum (with 0.02 % NaN_3_) (Invitrogen), 3 % donkey serum (Millipore) and 0.5 % Triton X-100 in PBS at RT for 3 h. To detect TMEM16A and TMEM16B, cryosections and / or brain slices were incubated at 4 °C for 24 h in primary antibody solution containing blocking solution and a rabbit polyclonal antibody directed against TMEM16A (1:500) (Heinze, Seniuk et al. 2013) and a guinea pig polyclonal antibody directed against TMEM16B (1:100) (Billig, Pal et al. 2011). To visualize blood vessels in brain slices, sections were co-stained with a primary monoclonal rat antibody against CD31 (1:500, BioLegend (102502)). Cryosections of the cochlea and brain slices were washed in PBS for 1 h and subsequently incubated in secondary antibody solution containing blocking solution and the secondary antibodies Alexa Fluor 488-conjugated goat anti-rabbit (1:1000, cochlear cryosections) (Molecular Probes (A11008)), Cy3-conjugated donkey anti-rabbit (1:1000, brain slices) (Jackson Immuno Research (711-165-152)) and DyLight 488-conjugated goat anti-rat (1:1000, brain slices) (Bethyl laboratories (A110-305D2)) at 4 °C for more than 12 h. Cochlear cryosections and brain slices were washed with PBS for 1 h, counterstained with DAPI to visualize cell nuclei and mounted on microscope slides (ProLong Gold anti fade reagent, Life Technologies). Images were captured using an upright confocal microscope (LSM 710, Zeiss).

### Hematoxylin-Eosin staining

Cochlear cryosections (see immunohistochemistry) were fixed in 4 % PFA (in PBS) at RT for 20 min. For Hematoxylin-Eosin (HE) staining, the Robot-Stainer device from Microm (HMS 740) was used.

### Western blot

Modioli including the SGNs were isolated from *Tmem16b* knockout mice and their wildtype littermates (P5). For immunoblots the tank blotting Mini-Protean Tetra system (Bio-Rad) and transfer buffer with methanol (25 mM Tris, 192 mM glycine) were used. 10 μg of proteins were separated in 5 % acrylamide gels. Proteins were blotted on polyvinylidene membranes (Roti-PVDF, Roth). Protein transfer was controlled by staining with Ponceau S (0.2 % Ponceau S, 3 % acetic acid). Blocking was done in Tris buffered saline (TBS) supplemented with 0.05 % Tween-20 and 5 % dry milk powder. TMEM16B was detected using the same antibody as described in the immunohistochemistry section (1:1000). It was incubated overnight at 4 °C in TBS-T with 1 % dry milk powder. Washing was done with TBS / 0.05 % Tween-20. The secondary antibody (Goat anti-guinea pig IgG antibody-HRP, Merck, 1:1000) was incubated at RT for 2 h in TBS-T with 1 % dry milk powder. Signals were visualized by chemiluminescence (Amersham ECL Prime Western Blotting Detection Reagent (GE Healthcare). Documentation was done with the camera system Stella 3200 (Raytest) and the Xstella software.

### DIC / Ca^2+^ imaging

After isolation of the inner ear from 5-to-7-day-old mice, the cochlea was removed from the capsule and the spiral ligament was dissected away. The cochlea was then mounted on a poly-L-lysine-coated (100 μg / ml) petri dish. Dissections and imaging were done in artificial cerebrospinal fluid (ACSF) containing (in mM): 130 NaCl, 3.5 KCl, 2 CaCl_2_, 1.3 MgCl_2_, 1.2 NaHPO_4_, 25 NaHCO_3_ and 25 D-glucose (320 mOsm, pH 7.4). The ACSF was saturated with 95 % O_2_/5 % CO_2_.

For differential interference contrast (DIC) imaging, acutely isolated cochleae were transferred to an inverted microscope (Axio Observer.Z1, Zeiss). A timelapse image series was taken for 20 min with a 25 x water-immersion objective. One image per second was acquired (1392×1040 px) using a CCD camera (Photometrics Cool Snap HQ^2^, Visitron) and the Metamorph imaging software (Visitron).

Ca^2+^ signals were visualized by Ca^2+^ imaging of cochleae that had been bath-loaded with 10 μM Fura-2 AM (Invitrogen) and 0.02 % pluronic acid F-127 (Invitrogen) in ACSF at RT for 35-45 min. Loading was followed by another 30 min incubation in ACSF. Cochleae were imaged on an inverted microscope (Axio Observer.Z1, Zeiss) using a 25 x water-immersion objective. Using the MetaFluor imaging software (Visitron), 400 timelapse ratio images (1392×1040 px) were obtained at a rate of one image per second. Therefore, pairs of images were acquired at alternate excitation wavelengths (340 / 380 nm). The emission was filtered at 500-530 nm.

Images acquired during DIC- and Ca^2+^ imaging were processed using a custom written ImageJ macro and the ImageJ software (NIH). Each frame (time point tn) was subtracted from an average of five preceding frames: 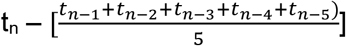. The mean areas and frequencies of volume changes and Ca^2+^ events were analyzed within a 10.000 μm^2^ area of Kölliker’s organ that was chosen from the center of the field of view. The frequencies of volume changes and Ca^2+^ events were analyzed by counting the number of events per second. The area of each event was measured in square micrometer using the Metamorph or MetaFluor imaging software for volume changes or Ca^2+^ signals, respectively. To be regarded as an event, volume changes / Ca^2+^ signals had to match the following two criteria: First, events must show an increase in fluorescence intensity / DIC contrast greater than 10 % of the baseline level. Baseline fluorescence / DIC contrast levels in an area of the cochlea that did not show volume changes or Ca^2+^ events (i.e. an area that lay outside of Kölliker’s organ and the IHC region). Second, changes in fluorescence / DIC contrast had to be larger than the surface area of one ISC. Videos were generated with the ImageJ software (7 images / s) and annotated using Premiere Pro (Adobe).

### *In vitro* electrophysiological recording and functional mapping

The brain was removed from mice (P9-11) and the forebrain and parts of the cerebellum were dissected to isolate the brainstem. Coronal brainstem slices (300-400 μm) were cut on a vibrating microtome (Leica VT1200S) in ice-cold dissection solution containing (in mM): 50 NaCl, 2.5 KCl, 1.25 NaH2PO_4_, 3 MgSO_4_, 0.1 CaCl_2_, 75 sucrose, 25 D-glucose, 25 NaHCO_3_ and 2 Na-pyruvate (280 mosmol, pH 7.2). The solution was oxygenated with 95 % O_2_ and 5 % CO_2_ during dissection. Slices recovered for 30 min at 36 °C and for an additional 30 min at RT in oxygenated recording solution containing (in mM): 125 NaCl, 2.5 KCl, 1.25 NaH_2_PO_4_, 2 MgSO_4_, 2.5 CaCl_2_, 18 D-glucose and 25 NaHCO_3_ (290 mosmol, pH 7.2). For electrophysiological recordings, brain slices were transferred to a recording chamber, mounted on an upright microscope (Axio Examiner A. 1, Zeiss), and perfused with oxygenated recording solution at a speed of 2 ml / min using a pressure driven perfusion system (ALA-VM8, ALA Instruments). The LSO was identified using a digital camera (C10600 Orca R^2^, Hamamatsu). Recordings were made from neurons with bipolar morphology from the medial (high frequency) region of the LSO in order to obtain comparable results and because the highest degree of refinement was reported for this frequency region (Sanes and Siverls, 1991). Electrodes had a resistance of 4-5 MΩ (Biomedical Instruments) when filled with intracellular solution containing (in mM): 54 potassium gluconate, 56 KCl, 1 MgCl2, 1 CaCl2, 5 sodium phosphocreatine, 10 HEPES, 11 EGTA, 0.3 Na-GTP and 2 Mg-ATP (285 mosmol, pH 7.2). Recordings were done in current clamp in the whole-cell configuration at RT and LSO neurons were held at −70 mV. Currents were acquired with an Axon CNS MultiClamp 700B amplifier (Molecular Devices) and low-pass filtered (3 kHz) and digitized (3 kHz) with a Digidata 1440A analog-to-digital converter (Molecular Devices) using the pClamp10 software (Molecular Devices).

### Photolysis of caged glutamate

Presynaptic MNTB neurons were localized by local uncaging of 120 μM MNI-caged glutamate trifluoroacetate (Femtonics) that was added to the recording solution. Uncaging of glutamate was controlled by a fast galvanometric photolysis system (UGA-40, Rapp OptoElectronic) that was triggered via a TTL-impulse. A 405 nm continuous diode laser (DL-405, Rapp OptoElectronic) was used as light source. Laser flashes were delivered through a light guide of 20 μm diameter (Rapp OptoElectronic) with a 10 x water immersion objective which produced circular spots of 20 μm diameters in the focal plane. For mapping, a virtual grid was defined over (and around) the MNTB harboring 250-300 grid points (spaced 20 μm apart) using the UGA-40 control 1.1 software (Rapp OptoElectronic). Each grid point represented one uncaging site. Laser pulses of 10 ms duration were delivered with 6 s delay time between uncaging sites. This resulted in reproducible LSO-MNTB input maps confirmed by rescanning in some cases. Only one MNTB map was obtained per mouse.

### Reconstruction and analysis of MNTB-LSO input maps

Events were detected with template search methods during a 20 ms window starting from the onset of the laser illumination. Peak amplitudes of the postsynaptic potentials (PSP) were measured (Clampfit 10.2 software, Molecular Devices). Each uncaging site that evoked PSP amplitudes greater than 2 mV in the recorded LSO neuron was considered as a functional MNTB-LSO connection and marked as colored square according to the following color code: yellow= depolarizations >2 mV, orange > 10 mV, red= action potential. Stimulation sites that evoked PSPs smaller than 2 mV or no PSPs were considered as unresponsive MNTB areas and were left blank. The total area covered by all colored squares was defined as the MNTB input area. The analysis of the MNTB input area was done in pixels (one pixel represents 1 μm^2^). The MNTB input area was calculated by multiplying the size of an uncaging site (~400 px) by the number of colored squares. The MNTB input area was normalized to the cross-sectional area of the MNTB. MNTB borders were determined by two investigators (one of them was blind to the mouse group) using images taken from the slices during the experiment. The result represents the MNTB input area in percent. The MNTB input width was calculated by measuring the maximal distance of stimulations sites that evoked depolarizations greater than 10 mV along the mediolateral axis. This distance was normalized to the mediolateral length of each MNTB. The result expresses the input width in percent along the tonotopic axis of the MNTB. On rare occasions, responses could be elicited from uncaging sites slightly ventral or dorsal to the defined MNTB boundaries in mice of both genotypes. These sites were included in the analysis.

### Spatial resolution of glutamate uncaging

The spatial resolution was assessed by direct current clamp whole cell recordings from MNTB neurons. The holding potential was −70 mV and the electrode resistance was 3-4 MΩ when filled with intracellular solution containing (in mM): 54 potassium gluconate, 56 KCl, 1 MgCl_2_, 1 CaCl_2_, 5 sodium phosphocreatine, 10 HEPES, 11 EGTA, 0.3 Na-GTP and 2 Mg-ATP (285 mosmol, pH 7.2). A virtual 10×10 grid was defined over the recorded MNTB neuron with uncaging points spaced 10 μm apart. The “action potential-eliciting distance” was defined as the maximal distance from the center of the uncaging site to the center of the cell body at which uncaging produced action potentials in the recorded MNTB neuron. The action potential-eliciting distance was measured for the mediolateral and the dorsoventral direction of the MNTB. We used the determined action potential-eliciting distance to define the uncaging parameters for the mapping experiment.

### *In vivo* recordings in the MNTB

Juxtacellular single unit recordings were performed in mice before or right after hearing onset (P8, P14). Animals were anesthetized with an initial intraperitoneal injection of a mixture of ketamine hydrochloride (0.1 mg / g body weight; Ketavet, Pfizer) and xylazine hydrochloride (5 μg / g body weight; Rompun, Bayer). Throughout recording sessions, anesthesia was maintained by additional subcutaneous application of one third of the initial dose, approximately every 90 min. The MNTB was approached dorsally and typically reached at penetrations depths of 5000-5500 μm.

The experimental protocol for acoustic stimulation and single unit recording was described in detail previously (Sonntag, Englitz et al. 2009, Dietz, Jovanovic et al. 2012). Briefly, auditory stimuli were digitally generated using custom-written Matlab software (The MathWorks Inc, Natick, USA) at a sampling rate of 97.7 kHz. The stimuli were transferred to a real-time processor (RP2.1, Tucker-Davis Technologies), D/A converted and delivered through custom-made earphones (acoustic transducer: DT 770 pro, Beyer Dynamics). Two recording protocols were used: (1) Pure tone pulses (100 ms duration, 5 ms rise-fall time, 100 ms interstimulus interval) were presented within a predefined matrix of frequency / intensity pairs to determine the excitatory response areas of single units. Characteristic frequency (CF, sound frequency causing a significant increase of unit’s action potential spiking at the lowest intensity), the respective threshold, Q-values (Qn, a measure of the unit’s sharpness of tuning calculated as the ratio of CF to bandwidth at n= 10, 20 and 30 dB above threshold), and maximum discharge rates were computed from the response area and used for the next protocol: (2) Spontaneous neuronal discharge activity was assessed in absence of acoustic stimulation to determine average firing rate and coefficient of variation (CV) of interspike intervals (ISIs = time that passes between two successive action potentials). Regularity of spontaneous discharge activity was quantified by CV, calculated as the ratio of standard deviation of ISIs and mean ISI. Additional analysis was done for MNTB units recorded in P8 mice, where spontaneous spiking discharges are grouped in bursts followed by periods of greatly reduced discharge activity (Sonntag et al., 2009). We used a statistical test based on gamma probability distribution of ISIs to determine the number and duration of bursts, and number of spikes per burst within each single cell recording.

For the juxtacellular single unit recordings glass micropipettes (GB150F-10, Science Products) were pulled on a horizontal puller (DMZ universal puller, Zeitz) to have a resistance of 5-10 MΩ when filled with 3 M KCl. Recorded voltage signals were amplified (Neuroprobe 1600, A-M Systems), bandpass filtered (0.3-7 kHz), digitized at a sampling rate of 97.7 kHz (RP2.1, Tucker-Davis Technologies), and stored for offline analysis using custom-written Matlab software. Three criteria were used to classify single unit recordings: (i) changes in the spike height did not exceed 20%, (ii) uniform waveforms, (iii) signal-to-noise ratio at least 8:1. Principle neurons of the MNTB were identified by the complex waveform of the recorded discharges (Guinan and Li 1990, Sonntag, Englitz et al. 2009).

Histological verification of the recording site was done by iontophoretic injection of Fluorogold (4 μA for 7 min). The animal was perfused 4-6 h after injection with 0.9 % NaCl solution followed by 5 % PFA. The brain was cut on a vibratome (Microm HM650) and the tissue sections (100 μm thick) were visualized under the fluorescent microscope (Axioskop, Zeiss). An examples of a recording site is shown in Figure 2I.

### Auditory brainstem response (ABR)

ABR recordings were conducted as described previously (Jing et al., 2013). In brief, mice (P13-14) were anesthetized intraperitoneally with a combination of ketamine (125 mg/kg) and xylazine (2.5 mg/kg). The ECG was constantly monitored to control the depth of anesthesia. The core temperature was maintained constant at 37° C using a temperature-controlled heat blanket (Hugo Sachs Elektronik–Harvard Apparatus). Note that in these small immature mice, the temperature probe could not be placed rectally, contributing to the relatively long latencies and greater variability of the ABR waves. ABR peaks IV (~5.3 ms) and V (~7ms), were very small and were thus excluded from analysis. For stimulus generation, presentation, and data acquisition, the TDT System II (Tucker Davis Technologies) was used that was run by the BioSig32 software (Tucker Davis Technologies). SPLs were provided in dB SPL root mean square (RMS) (tonal stimuli) or dB SPL peak equivalent (clicks) and were calibrated using a 1/4 inch Brüel and Kjær microphone (model 4939). Tone bursts (6/12/24 kHz, 10 ms plateau, 1 ms cos ^2^ rise / fall) or clicks of 0.03 ms were presented at 40 Hz or 20 Hz, respectively, in the free field ipsilaterally using a JBL 2402 speaker. The difference potential between vertex and mastoid subdermal needles was amplified (50.000x) and filtered (400-4000 Hz, NeuroAmp) and sampled at a rate of 50 kHz for 20 ms, 1300x to obtain two mean ABRs for each sound intensity. Hearing threshold was determined with 10 dB precision as the lowest stimulus intensity that evoked a reproducible response waveform in both traces by visual inspection.

### Statistics and statistical significance

For statistical analysis the SigmaPlot 12.5 and (Systat Software Inc) and GraphPadPrism software was used. Datasets with normal (Gaussian) distribution were assessed by unpaired Student’s t-tests (two-tailed distribution) if not said otherwise. For data sets with non-Gaussian distribution the non-parametric Mann-Whitney Rank Sum test was used. For analysis of the intensity-dependence of ABR amplitudes and latencies and of ABR thresholds across frequencies, 2way ANOVA was used. If the data were normally distributed the results are displayed as means ± standard error of means (s.e.m.) and otherwise as medians with the respective 25 % and 75 % quartiles. Mean cumulative distributions of inter-event intervals and distribution of ISIs between wildtype and cKO groups were compared using the two-sample Kolmogorov-Smirnov test and the chi-square test, respectively (see Figure 2). To avoid assigning too much weight on high rate-spiking cells, distributions were created by pooling n random ISIs for each cell within the two groups (n = lowest number of events recorded in any of the cells). A statistical test based on gamma probability distribution of ISIs was used to determine the number and duration of bursts and number of spikes per burst within each single cell recording. In brief, we assumed that action potential firing of MNTB neurons is a random Poisson-like process. In this way, the probability to encounter one ISI in time period τ is p = 1-e^-λτ^, where λ stands for neuronal firing rate. Next, the probability to encounter *k* ISIs during the time period τ as *k*-fold convolution of exponential distribution density was calculated, thereby yielding gamma distribution of waiting times for *k* ISIs. The resulting probability is 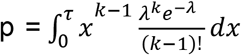. We ran the statistical test through our recorded action potential times and each spike train where probability stayed lower than 0.01 (p < 0.01) for at least 10 ISIs (k≥10) was recognized as a burst. Significance levels < 0.05 were assigned one star, < 0.01 two stars, and < 0.001 three stars. No statistical methods were used to predetermine sample sizes.

## Author contributions

A.M. performed and analyzed *in vitro* electrophysiological, immunohistochemical and Ca2+- /DIC imaging experiments, S.J. performed and analyzed *in vivo* electrophysiological experiments, N.S. performed and analyzed ABR experiments, A.K.H and C.A.H. generated the TMEM16A mouse and TMEM16A/B antibodies, T.M., R.R., and C.A.H. designed the experiments, A.M., S.J. and C.A.H. wrote the manuscript.

## Acknowledgements

We thank Andrew Groves for the opportunity to use the Pax2Cre mouse line. We further acknowledge technical support provided by Anje Sporbert and Zoltan Cseresnyes from the microscope core facility of the Max Delbrück Center for Molecular Medicine in the Helmholtz Association. This study was funded by a grant of the Thyssen foundation to Björn Schroeder and C.A.H. and by grants of the DFG to C.A.H. (HU 800/10-1) and to N.S., R.R. and T.M. (priority program 1608).

## Notes

### Competing Interest Statement

The authors have declared no competing interest.

