## Supplementary Figures and Tables for "The Cl^-^-channel TMEM16A controls the generation of cochlear Ca^2+^ waves and promotes the refinement of auditory brainstem networks"

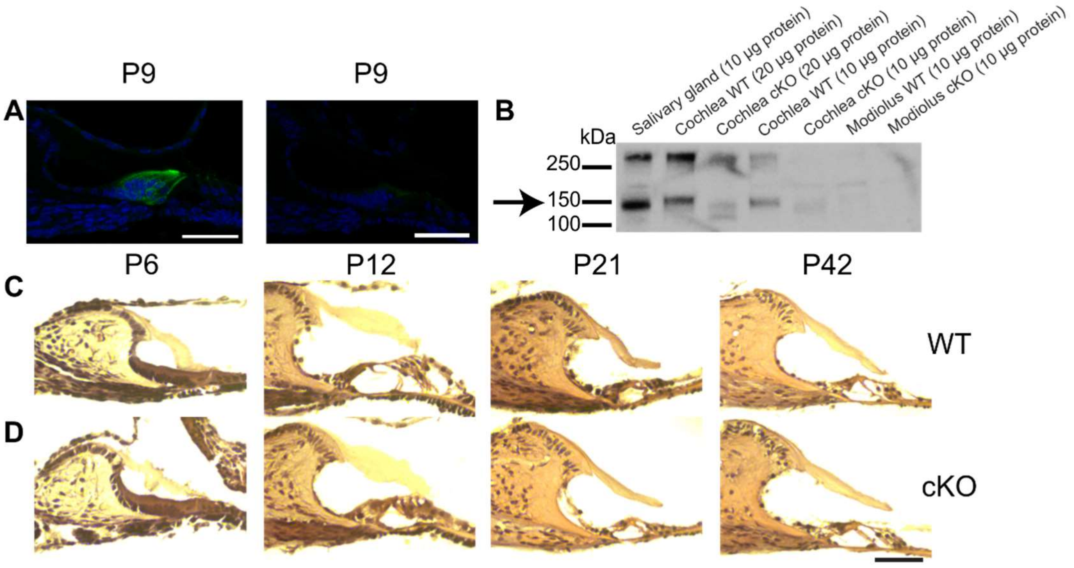


**Figure S1** Comparison of morphology and TMEM16A expression patterns in the developing organ of Corti between wildtype and cKO mice. (**A**) Immunohistochemical stainings for TMEM16A of cryosections from the organ of Corti in P9 wildtype (*left*) and cKO (*right*) mice. TMEM16A is broadly expressed in plasma membranes of ISCs and phalangeal cells in the wildtype organ of Corti, while the cKO cryosection shows a faint TMEM16A staining in the ISC area of Kölliker's organ, indicating that the TMEM16A antibody recognizes the truncated TMEM16A protein. Green - TMEM16A; blue - DAPI. Scale bar 50 µm. (**B**) Western blot analysis of protein lysates reveal that TMEM16A (*arrow*) is expressed in the salivary gland as previously reported (Romanenko et al. 2010), which is here used as positive control. The upper band represents TMEM16A dimers. TMEM16A is also expressed in the cochlea of wildtype mice. The TMEM16A signal in the cochlea migrates at a slightly higher molecular weight compared to the salivary gland, which might be explained by a difference in glycosylation or tissue specific splicing. In cKO mice the bands corresponding to TMEM16A were shifted to a smaller size confirming that the knockout was successful (Heinze et al., 2013). In contrast, no expression of TMEM16A is detected in the modiolus in wildtype and cKO mice. (**C, D**) HE stainings of organ of Corti cryosections dissected at different ages do not reveal morphological differences between wildtype and cKO mice between 1-6 weeks of age (P6, P12, P21, P42). Scale bar 50 µm.

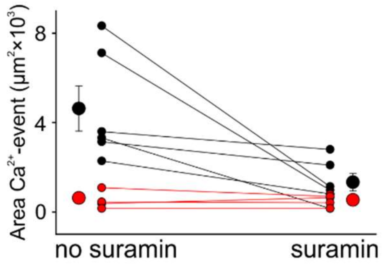


**Figure S2** Absence of P2 receptor activation in cKO mice. The purinergic receptor blocker suramin reduces $Ca^{2+}$-waves to uncoordinated $Ca^{2+}$ transients in wildtype (black) but does not have an effect on $Ca^{2+}$ transients in cKO littermates (red) (mean values ± SEM, n=6 WT, n=9 cKO (P5-7); two-tailed paired Student's t-test comparing area of $Ca^{2+}$ event before suramin application versus area of $Ca^{2+}$ event after suramin application: $p<0.05$ WT, $p=0.24$ cKO).

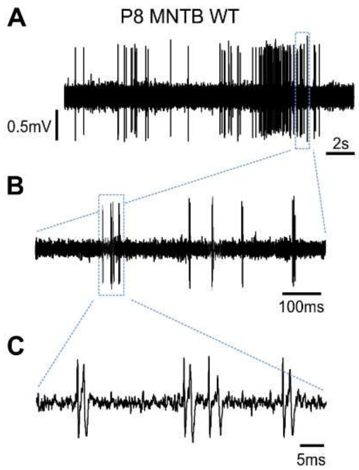


**Figure S3** Spontaneous activity of an exemplary P8 MNTB wildtype neuron *in vivo*. **(A)** A representative 20s voltage trace showing one prominent 3.5s burst. **(B)** Fine structure of the burst shown in A, reveals mini-bursts separated by 50-200ms silent periods. **(C)** The mini-burst shown in B is composed of four successive spikes. Note that each spike is composed of the prepotential, coming from the calyx of Held and the action potential from the MNTB principal neuron.

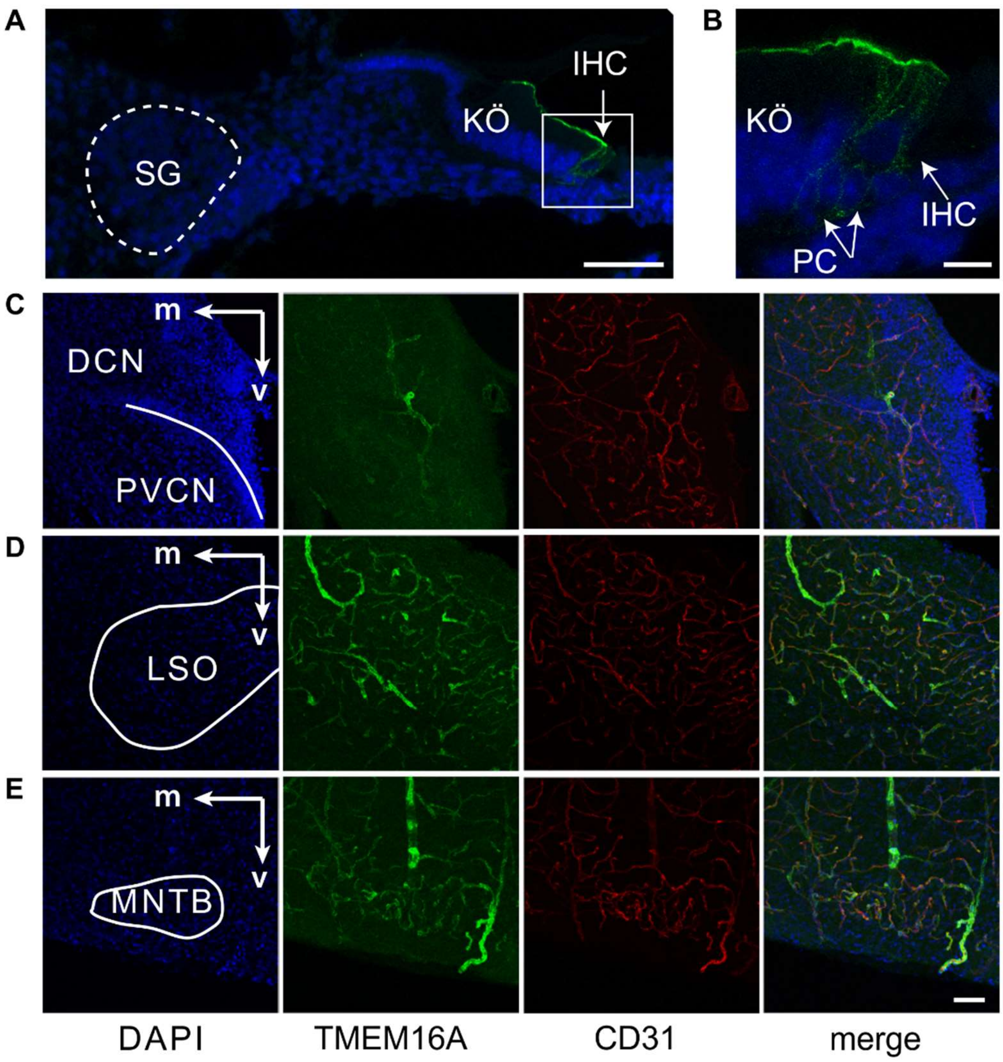


**Figure S4** TMEM16A is not expressed in SG, CN, MNTB and LSO neurons before hearing onset in wildtype mice. (**A**) At P2, TMEM16A is expressed in ISCs of Kölliker's organ and phalangeal cells but not in SG neurons (*dashed line*). IHC, inner hair cells; KÖ, Kölliker's organ; SG, spiral ganglion. Scale bar 50 µm. (**B**) Magnification of the IHC region from (A). PC, phalangeal cells. Scale bar 10 µm. (**C - E**) Immunohistochemical staining of brainstem slices for TMEM16A (*green*) and the endothelial marker CD31 (*red*) of wildtype mice (P8) showing TMEM16A expression in vascular smooth muscle cells (Heinze, Seniuk et al. 2013), but no expression in neurons of the CN (C), LSO (D), or MNTB (E). m, medial; v, ventral; DCN, dorsal cochlear nucleus; LSO, lateral superior olive; MNTB, medial nucleus of the trapezoid body; PVCN, posteroventral cochlear nucleus. Scale bars 50 µm.

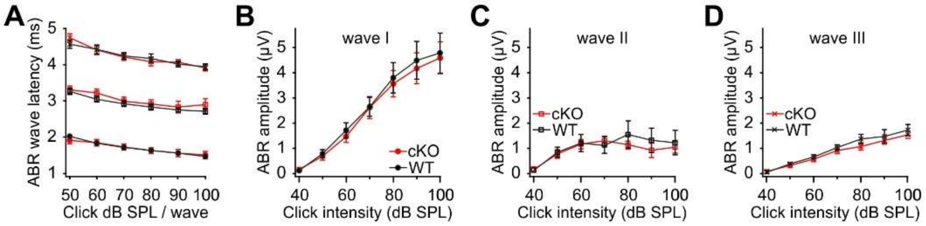


**Figure S5** Wildtype and cKO mice have similar auditory brainstem responses (ABRs). Shown are latencies (**A**) and amplitudes (**B** - **D**) of the first three ABR peaks (wave I – III) in response to click stimuli presented at different intensities (40 to 100 dB) (n=6 WT; n=7 cKO; P13-14).

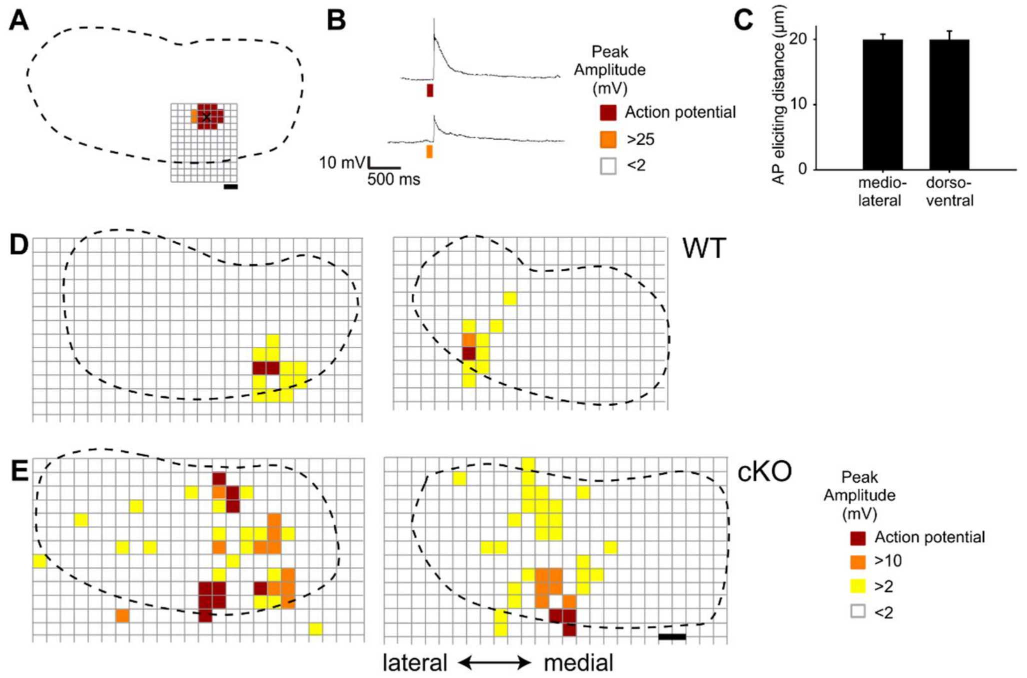


**Figure S6** Spatial resolution of glutamate uncaging and additional examples of MNTB input maps recorded from wildtype and cKO mice (**A**) Schematic representation of the MNTB from a wildtype mouse recorded at P8 (*dotted line* represents the MNTB area, *grid points* show glutamate uncaging sites, *black cross* indicates the location of the recorded MNTB neuron). Spatial resolution of glutamate uncaging was assessed by whole-cell current clamp recordings. Stimulation sites that evoked action potentials (*red squares*) were limited to the close proximity of the recorded MNTB neuron. Stimulation sites that were located in greater distance to the recorded MNTB neuron evoked subthreshold responses (*orange squares*) or no response (*white squares*). Scale bar 20 µm. (**B**) Uncaging of glutamate close to the recording site strongly depolarizes or elicits action potentials in the recorded MNTB neuron (see color code). (**C**) Quantification of the maximal action potential eliciting distance from the uncaging site to the center of recorded MNTB neuron. Values represent mean ± SEM (n=6 (P8-11)). (**D**) Two MNTB-LSO input maps from wildtype mice (P11, left; P9, right). (**E**) Two MNTB-LSO input maps from two cKO mice (P10, left; P11, right). Scale bar 40 µm.

**Table S1.** Comparison of the distribution of ISIs between wildtype and cKO mice.

| ISI | mean No ISIs (%) WT | mean No ISIs (%) cKO | p-value Chi Square test |
|---|---|---|---|
| **0 - 1.0** | 0.00 | 0.08 | 0.3340 |
| **1.0 - 1.58** | 0.09 | 0.00 | 0.3006 |
| **1.58 - 3.98** | 12.05 | 12.45 | 0.7785 |
| **3.98 - 6.31** | 14.72 | 13.17 | 0.3105 |
| **6.31 - 10** | 9.12 | 7.47 | 0.1589 |
| **10 - 15.85** | 6.88 | 2.41 | 2.8664E-07 |
| **15.85 - 25.12** | 6.28 | 2.65 | 2.2086E-05 |
| **25.12 - 39.81** | 7.23 | 2.97 | 1.0653E-07 |
| **39.81 - 63.1** | 6.80 | 4.82 | 0.1347 |
| **63.1 - 100** | 7.92 | 7.63 | 0.8008 |
| **100 - 158.5** | 10.41 | 11.49 | 0.4271 |
| **158.5 - 251.2** | 6.37 | 11.49 | 2.9338E-05 |
| **251.2 - 398.1** | 3.87 | 8.51 | 5.5438E-06 |
| **398.1 - 631** | 3.18 | 4.82 | 0.0458 |
| **631 - 1000** | 2.24 | 2.89 | 0.3178 |
| **1000 - 1585** | 0.95 | 2.17 | 0.0287 |
| **1585 - 2512** | 0.77 | 1.93 | 8.4155E-03 |
| **2512 - 3981** | 0.69 | 0.72 | 0.9200 |
| **3981 - 6310** | 0.26 | 1.12 | 0.0115 |
| **6310 - 10000** | 0.17 | 1.20 | 2.5904E-03 |

**Table S2.** Quantification of ABR thresholds (mean ± SEM) in response to stimulation with tone-bursts at 6 kHz, 12 kHz, and 24 kHz, or click stimulation.

| tone frequency (kHz) | mean ABR threshold (dB SPL) WT | mean ABR threshold (dB SPL) cKO | p-value |
|---|---|---|---|
| **6 kHz** | 60.0 ± 5.7 | 63.3 ± 3.4 | 0.53, 2way ANOVA |
| **12 kHz** | 43.3 ± 3.7 | 41.4 ± 2.8 | |
| **24 kHz** | 49.2 ± 3.0 | 54.3 ± 5.3 | |
| **click** | 45.0 ± 2.5 | 46.4 ± 3.0 | 0.91 (Mann-Whitney) |

**Table S3.** Quantification of peak amplitudes (mean ± SEM) of the first three ABR waves (I-III) in response to click stimuli of various intensities (40–100 dB).

| wave | click intensity (dB SPL) | mean amplitude WT | mean amplitude cKO | p-value, 2way ANOVA |
|---|---|---|---|---|
| **I** | **50** | 0.78 ± 0.16 | 0.68 ± 0.15 | 0.5175 |
| | **60** | 1.71 ± 0.30 | 1.47 ± 0.24 | |
| | **70** | 2.65 ± 0.38 | 2.59 ± 0.42 | |
| | **80** | 3.80 ± 0.60 | 3.56 ± 0.52 | |
| | **90** | 4.49 ± 0.75 | 4.17 ± 0.61 | |
| | **100** | 4.77 ± 0.80 | 4.59 ± 0.63 | |
| **II** | **50** | 0.77 ± 0.24 | 0.76 ± 0.16 | 0.3879 |
| | **60** | 1.30 ± 0.29 | 1.16 ± 0.18 | |
| | **70** | 1.24 ± 0.28 | 1.30 ± 0.17 | |
| | **80** | 1.66 ± 0.49 | 1.12 ± 0.16 | |
| | **90** | 1.43 ± 0.44 | 0.92 ± 0.29 | |
| | **100** | 1.37 ± 0.43 | 1.04 ± 0.23 | |
| **III** | **50** | 0.37 ± 0.04 | 0.31 ± 0.09 | 0.0807 |
| | **60** | 0.64 ± 0.07 | 0.45 ± 0.13 | |
| | **70** | 1.02 ± 0.11 | 0.77 ± 0.19 | |
| | **80** | 1.36 ± 0.18 | 0.89 ± 0.21 | |
| | **90** | 1.46 ± 0.27 | 1.09 ± 0.25 | |
| | **100** | 1.71 ± 0.22 | 1.27 ± 0.27 | |

**Table S4.** Quantification of latencies (mean ± SEM) of the first three ABR waves (I – III) in response to click stimuli of various intensities (40–100 dB).

| wave | click intensity (dB SPL) | mean latency (ms) WT | mean latency (ms) cKO | p-value 2way ANOVA |
|---|---|---|---|---|
| I | 50 | 2.02 ± 0.03 | 2.02 ± 0.03 | 0.9073 |
| | 60 | 1.83 ± 0.03 | 1.83 ± 0.03 | |
| | 70 | 1.71 ± 0.04 | 1.71 ± 0.04 | |
| | 80 | 1.64 ± 0.04 | 1.64 ± 0.04 | |
| | 90 | 1.55 ± 0.03 | 1.55 ± 0.03 | |
| | 100 | 1.47 ± 0.03 | 1.47 ± 0.03 | |
| II | 50 | 3.26 ± 0.067 | 3.30 ± 0.11 | 0.1225 |
| | 60 | 3.05 ± 0.07 | 3.21 ± 0.11 | |
| | 70 | 2.02 ± 0.07 | 2.99 ± 0.11 | |
| | 80 | 2.83 ± 0.09 | 2.91 ± 0.12 | |
| | 90 | 2.75 ± 0.06 | 2.83 ± 0.16 | |
| | 100 | 2.71 ± 0.08 | 2.89 ± 0.17 | |
| III | 50 | 4.56 ± 0.11 | 4.74 ± 0.19 | 0.9147 |
| | 60 | 4.42 ± .012 | 4.40 ± 0.14 | |
| | 70 | 4.24 ± 0.09 | 4.20 ± 0.09 | |
| | 80 | 4.17 ± 0.12 | 4.08 ± 0.09 | |
| | 90 | 4.01 ± 0.07 | 4.06 ± 0.05 | |
| | 100 | 3.94 ± 0.08 | 3.90 ± 0.06 | |
